# Altered hematopoietic system and self-tolerance in Bardet-Biedl Syndrome

**DOI:** 10.1101/2020.02.24.962886

**Authors:** Oksana Tsyklauri, Veronika Niederlova, Elizabeth Forsythe, Ales Drobek, Avishek Prasai, Kathryn Sparks, Zdenek Trachtulec, Philip Beales, Martina Huranova, Ondrej Stepanek

## Abstract

Bardet-Biedl Syndrome (BBS) is a pleiotropic genetic disease caused by dysfunction of primary cilia. The immune system of patients with BBS or another ciliopathy has not been investigated, most likely because hematopoietic cells do not form cilia. However, there are multiple indications that the impairment of the processes typically associated with cilia might influence the hematopoietic compartment and immunity. In this study, we analyzed clinical data of BBS patients as well as a corresponding mouse model of BBS4 deficiency. We uncovered that BBS patients have higher incidence of certain autoimmune diseases. BBS patients and animal models have elevated white blood cell levels and altered red blood cell and platelet compartments. Moreover, we observed that BBS4 deficiency alters the development and homeostasis of B cells in mice. Some of the hematopoietic system alterations were caused by the BBS-induced obesity. Overall, our study reveals a connection between a ciliopathy and the alterations of the immune system and the hematopoietic compartment.

## Introduction

Bardet-Biedl Syndrome (BBS) is a recessive genetic disorder caused by complete or partial loss-of-function mutations in any of more than 20 BBS genes known to date. BBS belongs to a group of ciliopathies, i.e., disorders caused by defective formation and/or function of primary cilia. Eight of the BBS proteins (BBS1, BBS2, BBS4, BBS5, BBS7, BBS8, BBS9, and BBS18) form a transport complex called the BBSome, which sorts selected cargoes into and out of the cilium [1–4]. Other commonly mutated BBS genes (*ARL6*/*BBS3*, *MKKS*/*BBS6*, *BBS10*, and *BBS12*) assist the BBSome assembly or function [1, 5]. The BBSome is believed to act as a cargo adaptor connecting the cargoes to the intraflagellar transport (IFT) machinery [6, 7].

BBS is a pleiotropic disease with rod-cone dystrophy, polydactyly, obesity, learning difficulties, hypogonadism, and renal anomalies being the primary diagnostic features [8]. The immune system of patients with ciliopathies including BBS has not been studied in detail. An exceptional study in this respect is a case report of 3 BBS patients suffering from autoimmune diseases in a cohort of 15 studied BBS patients [9]. Similarly, the immune system has not been thoroughly investigated in animal models of ciliopathies either. The possible connection between ciliopathies and the immune system has not been addressed most likely because immune cells do not form primary cilia [10, 11]. However, there are several lines of evidence suggesting that the BBS might affect the function of the immune system.

First, the immunological synapse formed between T cells and antigen-presenting cells exhibits a striking analogy to the primary cilium [12, 13]. Formation of both the immunological synapse and the cilium involves the reorganization of cortical actin and the centrosome polarization. Along this line, some components of the IFT machinery have been shown to participate in the organization of the immunological synapse to promote T-cell activation [14, 15]. In particular, it has been shown that the vesicles containing key T-cell signaling molecules TCR/CD3 complex and LAT are transported towards the immunological synapse by IFT proteins [16, 17].

Second, the BBSome is required for Sonic hedgehog (SHH) signaling [18–20]. The SHH signaling pathway regulates multiple processes in the organism including T-cell development [21]. In the thymus, SHH regulates the development of thymocytes before and soon after the pre–TCR signaling [22–24]. In the periphery, SHH has been shown to negatively regulate TCR-dependent differentiation of T cells [25], as well as to promote Th2 differentiation and allergic reactions [26]. Key components of the SHH signaling pathway, SMO, IHH, GLI1, and PTCH2 are upregulated in effector cytotoxic T cells and transported towards the immunological synapse in vesicles [27]. Moreover, *Smo*^KO/KO^T cells showed reduced cytotoxity associated with defects in actin remodeling required for the centrosome polarization and the release of cytotoxic granules [27].

Third, the BBSome regulates trafficking of the leptin receptor [28]. Leptin is a signaling molecule which acts as a pro-inflammatory cytokine [29, 30]. In particular, leptin signaling inhibits the proliferation of regulatory T cells [31] and promotes the effector cell proliferation and polarization towards Th1 helper T cells [32]. Moreover, T cells deficient in the leptin receptor show impaired differentiation into Th17 helper T cells in mice [33], indicating a T-cell intrinsic role of leptin signaling.

Fourth, one of the major symptoms of BBS is obesity, which is believed to undermine the immune tolerance [34]. Obesity induces production of pro-inflammatory cytokines, such as TNF-α [35] and IL-6 [36], which might predispose the individual for the development of autoimmune diseases [37–41]. Thus, the BBSome might have an extrinsic role in the immune system via inducing obesity.

In this study, we addressed the intrinsic and extrinsic roles of the BBSome in the immune system by investigating BBS patients and a BBS mouse model of the BBS4 deficiency. We uncovered that BBS patients show elevated prevalence of particular autoimmune diseases. We identified dysregulated homeostasis of blood cells both in BBS patients and in BBS4-deficient mice. Besides, we revealed the association of the BBS induced obesity with specific hematopoietic system alterations in BBS patients.

## Methods

### Antibodies and reagents

Antibodies to the following antigens were used for flow cytometry: CD4 BV650 (RM4-5, #100545, Biolegend), CD8a PE-Cy7 (53-6.7, #1103610, SONY), CD8a FITC (53-6.7, #100706, Biolegend), CD19 PE (6D5, #115508, Biolegend), CD23 APC (b3b4, #1108095, SONY), CD44 PE (IM7, #103008, Biolegend), B220 Alexa Fluor 700 (RA3-6B2, #103231, Biolegend), B220 FITC (RA3-6B2, #103206, Biolegend), CD69 PE (H1.2F3, #104508, Biolegend), IgM BV421 (rmm-1, #2632585, SONY), IgD Per-CP-Cy5.5 (11-26c.2a, #2628545, SONY), IgLλ APC (RML-42, #407306, Biolegend), TCRβ APC (H57-597, #109212, Biolegend).

Antibodies used for immunoblot analysis: BBS4 (rabbit, a kind gift from Prof. Maxence Nachury, UCSF, CA, USA), β-actin (mouse, #4967, Cell Signaling), α-mouse-HRP, α-rabbit-HRP (both from Jackson ImmunoResearch).

Antibodies used for lymphocyte enrichment: biotinylated α-TCRβ (H57-597, #553169, BD Pharmingen), α-CD19 (6D5, #115503, Biolegend).

4-hydroxy-3-nitrophenylacetic acid succinimide ester (LGC, Biosearch Technologies)

Peptides OVA (SIINFEKL), Q4R7 (SIIRFERL), Q4H7 (SIIRFEHL), T4 (SIITFEKL) were purchased from Eurogentec or Peptides&Elephants.

Dyes: CFSE and DDAO cell tracker dyes (both Invitrogen), LIVE/DEAD near-IR dye (Life Technologies), Hoechst 33258 (Life Technologies).

### Mice

All mice were 5-25 weeks old and had C57Bl/6J background. B1-8 [42], RIP.OVA [43], OT-I *Rag2*^KO/KO^ [44], *Vav-iCre* [45, 46], *Cd4-Cre* [47] strains were described previously. *Bbs4*^+/GT^ sperm (*Bbs4*^tm1a(EUCOMM)Hmgu^) was obtained from KOMP (UC Davis, CA, USA) and used for *in vitro* fertilization. *Bbs4*^+/+^ and *Bbs4*^GT/GT^ or *Bbs4*^KO/KO^ littermates were generated by intercrossing heterozygous animals. Mice were bred in specific-pathogen-free facility (Institute of Molecular Genetics) [48]. Animal protocols were approved by the Czech Academy of Sciences, in accordance with the laws of the Czech Republic.

Males and females were used for the experiments. If possible, age- and sex-matched pairs of animals were used in the experimental groups. If possible, littermates were equally divided into the experimental groups.

No randomization was performed since the experimental groups were based solely on the genotype of the mice. The experiments were not blinded since no subjective scoring method was used.

### qPCR of BBS genes in immune organs and T cells

Total RNA (1 or 2 μg) of organs (kidney, brain, lymph nodes, spleen) and T cells from C57BL/6J WT mice was obtained in 3 independent biological replicates and transcribed using RevertAid reverse transcriptase (Thermofisher, #EP0442) with oligo(dT)_18_ primers according to the manufacturer’s instructions. RT-quantitative PCR was carried out using LightCycler 480 SYBR green I master chemistry (Roche). All samples were measured in triplicates. Obtained C_T_ values were normalized to data of reference genes Glyceraldehyde-3-Phosphate Dehydrogenase (*Gapdh*), Tubulin Beta 2A Class IIa (*Tubb2a*) and Eukaryotic Translation Elongation Factor 1 Alpha 1 (*Eef1a1*). The sequences of used primers are

GAPDH: F TGCACCACCAACTGCTTAGC, R GGCATGGACTGTGGTCATGAG; Tubb2A: F
AACCAGATCGGCGCTAAGT, R TGCCAGCAGCTTCATTGTA; eEF1a1: F
ACACGTAGATTCCGGCAAGT, R AGGAGCCCTTTCCCATCTC; BBS1: F
ATCGGATTCTGACAGCGGG, R CCACCAGCTTGTACTCCCCA; BBS2: F
TGCCCCGATTCACCATGTAT, R CACGTGACCATCCTCTGTGTG; BBS4: F
AGCTTGGGATGAAAACTCAGGT, R GCTGTTCTTTGATCACAGCCTT; BBS5: F
GCGACCAGGGGAATTTAGGA, R ATGACAAGCGCCAAACCAAA; BBS7: F
AGGGCTACACAAAAGGTGGT, R TTCTCCTGAGGCGTGTTGAC; BBS8: F
CTTATGATCAGGCGGCTTGGA, R GTGGGACCTGAGCAATAGCA; BBS9: F
ACTCCAGACCGACAGGTATT, R GGCTGACCAGGTAGGCAAAT; BBip10: F
AGCCCCTGATCGCTTACCTA, R GACAATGTCTCACTCGTCAGC.

### Immunoblotting

Freshly isolated murine organs (testicles, thymi, brains) or enriched lymphocytes were homogenized in Laemmli sample buffer. The resulting lysates were separated on a polyacrylamide gel and transferred to nitrocellulose membrane using standard immunoblotting protocols. Membranes were probed with antibodies against BBS4 followed by secondary α-rabbit-HRP antibody. As a loading control we probed the membranes for β-actin followed by secondary α-mouse-HRP antibody. The images were obtained using chemiluminescence immunoblot imaging system Azure c300 (Azure Biosystems, Inc.).

### Histological analysis

Testes isolated from 30-day-old male mice were collected, immediately dipped into Bouin solution and fixed for 24 h at 4°C. Paraffin-embedded tissue blocks were cut with a microtome (Leica RM2255), and the sections were stained with hematoxylin/eosin using standard technique. The images were taken using microscope system Axioplan 2 imaging (Zeiss) using 10×/0.50 NA objective.

### Weighting of mice

Bodyweight of *Bbs4*^+/+^, *Bbs4*^GT/GT^, *Bbs4*^KO/KO^ mice was recorded weekly starting at 5 weeks of age. All the mice were kept in sex-matched cages together with their littermates (≤ 6 per cage), and fed a standard chow diet ad libitum.

### ELISA

Blood from *Bbs4*^KO/KO^, *Bbs4*^GT/GT^ and their age/sex-matched controls was collected by submandibular bleeding [49] into EDTA-coated tubes and centrifuged for 15 minutes at 1000 × g at 4°C in order to separate plasma. Obtained plasma samples were assayed immediately or stored at −80°C for later use. Leptin concentration was measured by mouse leptin ELISA Kit (Cloud-Clone Corp., SEA084Mu) according to the manufacturer’s instructions.

### Flow cytometry

Live cells were stained with relevant antibodies on ice. LIVE/DEAD near-IR dye or Hoechst 33258 were used for discrimination of live and dead cells. Flow cytometry was carried out using an LSRII (BD Bioscience). Data were analyzed using FlowJo software (TreeStar).

### B-cell activation

T2-Kb cells [50] were loaded with 4-hydroxy-3-nitrophenylacetic acid succinimide ester (NP-Osu) in PBS for 10 min at 37°C, washed and resuspended in RPMI/10% FCS. NP-loaded T2-Kb cells were mixed with splenocytes isolated from B1-8 mice (*Bbs4*^+/+^ and *Bbs4*^GT/GT^) at 1:10 or 1:3 ratios, and incubated for 6 hours at 37°C. After incubation, cells were centrifuged (1000 × g, 2 min), resuspended in PBS/0.5% gelatin, stained with antibodies (B220, IgLλ, CD69) for 30 min on ice, and analyzed by flow cytometry.

### T-cell conjugation assay

T-cell conjugation assay was performed as previously shown [44]. Briefly, OT-I T cells from *Bbs4*^FL/FL^ *Cd4-Cre*^−^ or *Bbs4*^FL/FL^ *Cd4-Cre*^+^ (cKO) littermates were stained with CFSE cell tracker dye, and splenocytes isolated from C57Bl/6 mice were stained with DDAO cell tracker dye. Splenocytes were loaded with OVA peptide or with indicated altered peptide ligands for 3 h in RPMI/10% FCS, mixed with OT-I T cells at 2:1 ratio, and centrifuged (1000 × g, 1 min). After 20 min of co-culture at 37°C/CO_2_ incubator, cells were fixed by adding formaldehyde (2% final, 35 min). Cells were centrifuged (1000 × g, 2 min), resuspended in PBS/0.5% gelatin, and analyzed by flow cytometry. Each of the four experiments was carried out in technical duplicates.

### Model of autoimmune diabetes

The model of autoimmune diabetes has been described previously [51]. Briefly, OT-I cells from *Bbs4*^FL/FL^ *Cd4-Cre*^−^ or *Bbs4*^FL/FL^ *Cd4-Cre*^+^ (cKO) sex-matched littermates were adoptively transferred into a host RIP.OVA mice intravenously. On the following day, the host mice were immunized with 5000 CFU of OVA expressing *Listeria monocytogenes* (Lm). Lm strain expressing OVA has been described previously [52]. Level of glucose in the urine of RIP.OVA mice was monitored on a daily basis using test strips (GLUKOPHAN, Erba Lachema).

The animal was considered to suffer from diabetes when the concentration of glucose in the urine reached ≥ 1000 mg/dl for 2 consecutive days. On day 7 post-infection, blood glucose was measured using contour blood glucose meter (Bayer).

### Blood analysis

Blood from 20-21 weeks old *Bbs4*^+/+^, *Bbs4*^KO/KO^ and *Bbs4*^GT/GT^ mice was collected by submandibular bleeding [49] into EDTA-coated tubes and analyzed using BC5300 Vet Auto Hematology Analyzer (Mindray Bio-Medical Electronics Co., Ltd.).

### Analysis of the clinical data of BBS patients

Fully anonymized medical records of 255 BBS patients were obtained from the Clinical Registry Investigating BBS (CRIBBS) by the NIH through the National Center for Advancing Translational Sciences and the Office of Rare Diseases Research (https://grdr.hms.harvard.edu/transmart). Data about the prevalence of autoimmune diseases in the CRIBBS cohort were compared to normal prevalence of autoimmune diseases reported in the Autoimmune Registry [53].

Medical records of BBS patients attending the BBS multidisciplinary clinic at Guy’s Hospital of Guy’s and St Thomas’ NHS Foundation Trust, London, or Great Ormond Street Hospital, London, were studied in detail with focus on presence of any immune-related phenotype. In addition to the manual control, the records were also automatically searched for the occurrence of the following terms: autoimm-, immun-, thyro-, inflam-, diabet-, T1DM, ulcerative, crohn, IBD, rheuma-, arthri-, joints. Statistical significance of the difference in the prevalence between the BBS patients and overall population was tested using two-tailed binomial test in RStudio (function binom.test).

Results of blood tests of BBS patients (total white blood cell count, leukocyte populations, hemoglobin, platelet counts, mean corpuscular volume, red blood cell count, hematocrit, red cell distribution width and mean corpuscular hemoglobin), their age ranges and body mass indices (BMI) were retrospectively ascertained from medical records stored at the BBS multidisciplinary clinic at Guy’s Hospital of Guy’s and St Thomas’ NHS Foundation Trust, London, or Great Ormond Street Hospital, London. Blood tests were performed during regular medical examination of the patients. All patients gave informed consent or assent. The protocol for this study was approved by the Great Ormond Street Hospital Research Ethics Committee (Project Molecular Genetics of Human Birth Defects – mapping and gene identification, reference #08/H0713/82) the and by the ethical committee of the Institute of Molecular Genetics of the ASCR.

Two distinct sets of controls for the analyzed set of BBS patients were selected from the 14750 participants of the UK Biobank project (ID: 40103) [54]. First, we selected 10 controls for each patient matching by age range (categories 41-50, 51-60, 60+ years) and sex. These controls had random BMI and thus were used as BMI-random controls. Second, we selected 10 controls for each patient matching by age range (categories 41-50, 51-60, 60+ years), sex, and BMI. These were used as BMI-matched controls. For 34 of the 42 patients we found controls with BMI difference ≤ 0.6 kg/m^2^. For the 8 patients with extreme BMI values, that precise matching was not possible, so that we selected the best-matching controls available for these cases. As the UK Biobank only includes participants older than 40 years, our analysis was limited to this age group.

### Enrichment of T and B lymphocytes

T and B lymphocytes were enriched by positive selection using the Dynabeads Biotin Binder kit (Invitrogen, #11047), and biotinylated α-TCRβ and α-CD19 antibodies, respectively.

## Results

### Autoimmune diseases are more prevalent in BBS patients

In this work, we studied the potential role of the BBSome in the immune system. Initially, we analyzed two cohorts of BBS patients from the CRIBBS NIH registry and from the Guy’s Hospital of Guy’s and St Thomas’ NHS Foundation Trust, London, or Great Ormond Street Hospital, London. We found out that certain autoimmune and inflammatory diseases, such as type I diabetes, Hashimoto’s thyroiditis, rheumatoid arthritis, and inflammatory bowel diseases, are more prevalent in BBS patients than in the overall population (Table I). These findings suggested that the BBSome has an intrinsic or extrinsic role in the immune system, particularly in the immune tolerance.

**Table 1.**
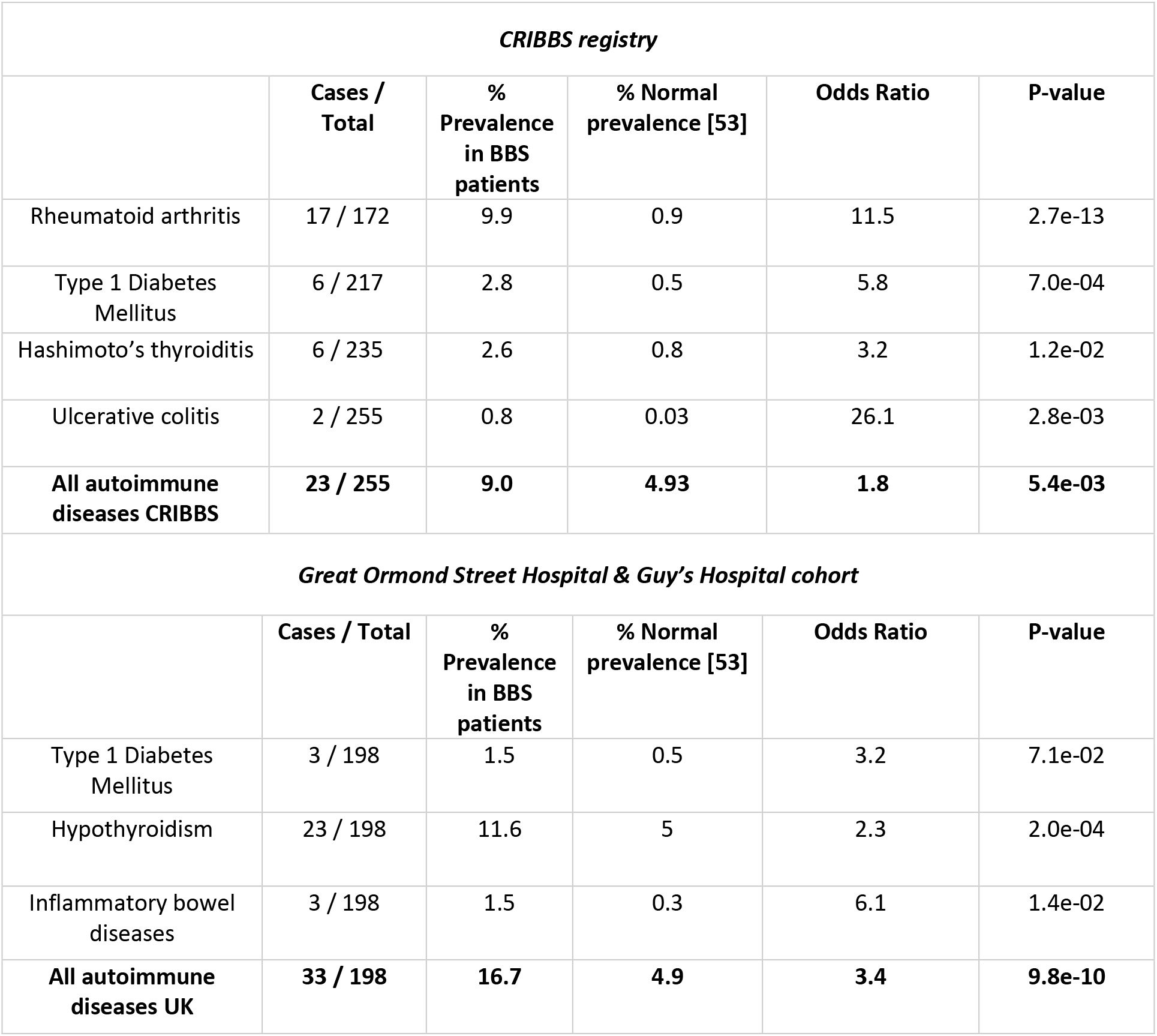
Autoimmune diseases in BBS patients. The table shows the fold change in the prevalence of autoimmune diseases in the CRIBBS cohort of 255 BBS patients (upper part) and in the cohort of 198 BBS patients from the Great Ormond Street Hospital and the Guy’s Hospital in London (lower part). Normal prevalence of autoimmune diseases was adopted from the Autoimmune Registry [53]. P value was calculated using binomial test.

| *CRIBBS registry* |  |  |  |  |  |
| --- | --- | --- | --- | --- | --- |
|  | **Cases / Total** | **% Prevalence in BBS patients** | **% Normal prevalence [53]** | **Odds Ratio** | **P-value** |
| Rheumatoid arthritis | 17 / 172 | 9.9 | 0.9 | 11.5 | 2.7e-13 |
| Type 1 Diabetes Mellitus | 6 / 217 | 2.8 | 0.5 | 5.8 | 7.0e-04 |
| Hashimoto's thyroiditis | 6 / 235 | 2.6 | 0.8 | 3.2 | 1.2e-02 |
| Ulcerative colitis | 2 / 255 | 0.8 | 0.03 | 26.1 | 2.8e-03 |
| **All autoimmune diseases CRIBBS** | **23 / 255** | **9.0** | **4.93** | **1.8** | **5.4e-03** |
| *Great Ormond Street Hospital & Guy's Hospital cohort* |  |  |  |  |  |
|  | **Cases / Total** | **% Prevalence in BBS patients** | **% Normal prevalence [53]** | **Odds Ratio** | **P-value** |
| Type 1 Diabetes Mellitus | 3 / 198 | 1.5 | 0.5 | 3.2 | 7.1e-02 |
| Hypothyroidism | 23 / 198 | 11.6 | 5 | 2.3 | 2.0e-04 |
| Inflammatory bowel diseases | 3 / 198 | 1.5 | 0.3 | 6.1 | 1.4e-02 |
| **All autoimmune diseases UK** | **33 / 198** | **16.7** | **4.9** | **3.4** | **9.8e-10** |

In the next step, we addressed the connection between the BBSome and the immune and hematopoietic systems using mouse models. First, we tested if the BBSome subunits are expressed in the murine immune tissues. We detected the expression of all 8 subunits in the spleen, lymph nodes, and isolated T cells on the mRNA level (Fig. 1A). The expression levels of *Bbs2, Bbs4, Bbs9*, and *Bbs18* in the lymphoid tissues were comparable to, or even slightly higher than in the brain and the kidney, two organs where the BBSome plays a major role [55–58]. The other four subunits (*Bbs1, Bbs5, Bbs7*, and *Bbs8*) were expressed in the lymphoid tissues at 10-to 50-fold lower levels than in the brain and the kidney. Moreover, we detected BBS4 protein in isolated T and B cells (Fig. 1B). Altogether, all the BBSome subunits are variably expressed in lymphocytes and lymphocyte-rich tissue, despite of the fact that hematopoietic cells are commonly considered as non-ciliated cells. This suggested that the BBSome as a whole or some individual BBSome subunits might have an intrinsic role in lymphocytes.

**Figure 1.**
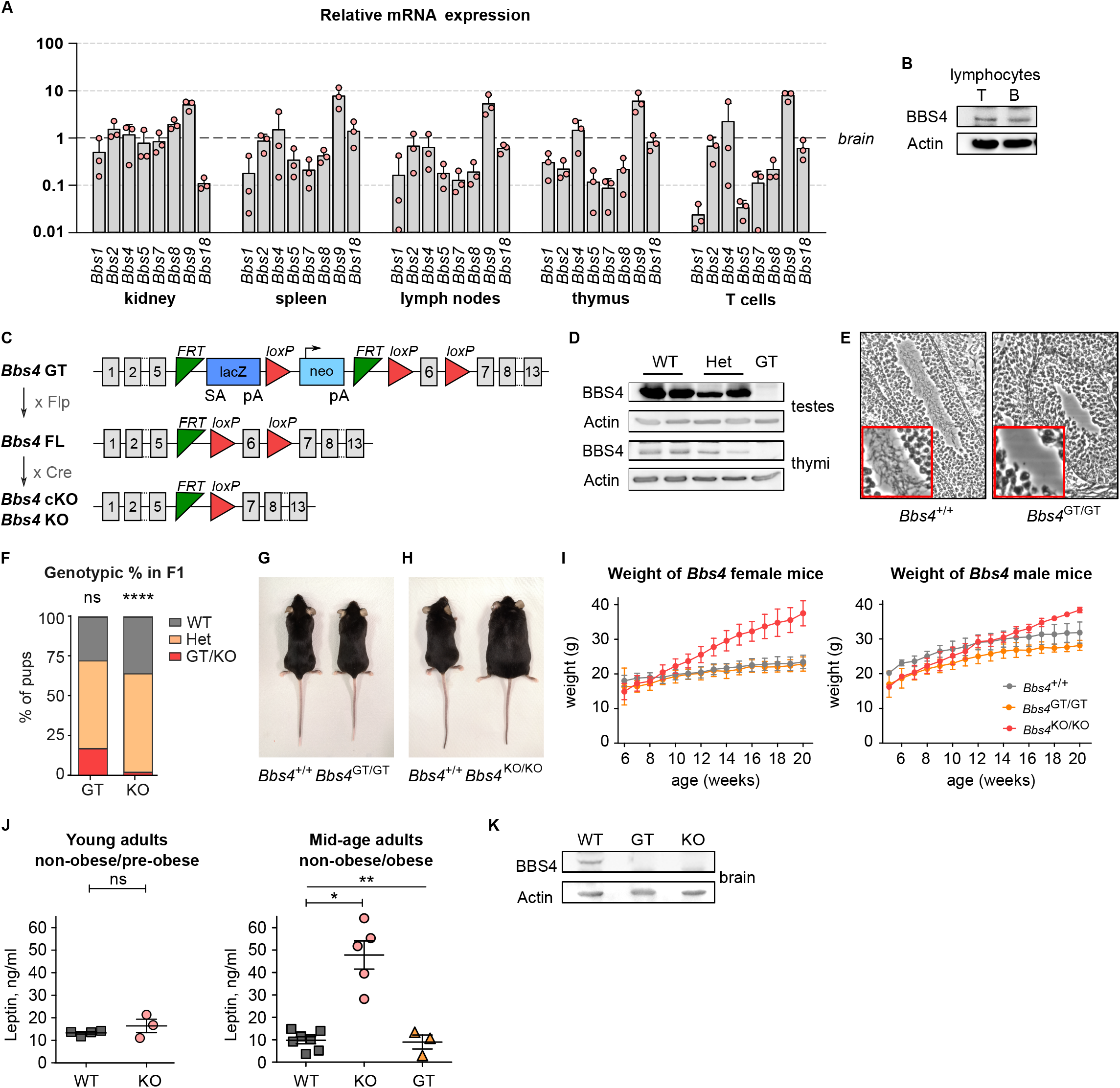
Mouse models for studying the BBSome in the immune system. **(A)** Relative expression of the BBSome subunits in the indicated organs and T cells measured by qPCR. C_T_ values of the BBS genes were normalized to the geometric mean of the C_T_ values of reference genes *Gapdh*, *Tubb2a* and *Eef1a1*. The expression levels are normalized to those of brain (=1). Mean+SD, 3 independent experiments. **(B)** Immunoblot analysis of BBS4 expression in T and B lymphocytes isolated from the lymph nodes and spleen of WT mouse. β-actin staining served as a loading control. A representative experiment out of 3 in total. **(C)** Schematic representation of mouse models of *Bbs4* deficiency used in this study. *Bbs4* GT, gene-trap allele interrupting the *Bbs4* gene. *Bbs4* FL, allele with floxed *Bbs4* exon 6. *Bbs4* KO and *Bbs4* cKO, alleles with deleted exon 6. **(D)** Immunoblot analysis of BBS4 expression in testicles and thymi lysates of WT, *Bbs4*^GT/+^ heterozygote and *Bbs4*^GT/GT^ mice. β-actin staining serves as a loading control. A representative experiment out of 3 in total is shown. **(E)** Hematoxylin and eosin staining of sections of seminiferous tubules from 30-days-old *Bbs4*^+/+^ and *Bbs4*^GT/GT^ males. A representative experiment out of 3 in total is shown. **(F)** Genotypic ratio of WT, heterozygous, or *Bbs4*-deficient offspring at weaning from mating of *Bbs4*^GT^/*+* (n=145 pups) or *Bbs4*^KO/+^ (n=168 pups) parents. Pearson’s chi-square (Χ^2^) test was used for statistical comparison of the observed distribution to the expected Mendelian ratio. **(G)** *Bbs4*^+/+^ and *Bbs4*^GT/GT^ female littermates at 20 weeks of age. Representative litter out of 7 in total. **(H)** *Bbs4*^+/+^ and *Bbs4*^KO/KO^ female littermates at 20 weeks of age. Representative litter out of 5 in total. **(I)** Growth curves of *Bbs4* deficient mice, mean ± SD is shown. Females: *Bbs4*^+/+^ (n=12), *Bbs4*^GT/GT^ (n=12), *Bbs4*^KO/KO^ (n=6). Males: *Bbs4*^+/+^ (n=4), *Bbs4*^GT/GT^ (n=7), *Bbs4*^KO/KO^ (n=3). **(J)** Leptin concentration in blood plasma taken from mid-age (14-20 weeks) adult mice, and young (7-8 weeks) adult mice. Young adult mice: *Bbs4*^+/+^ (n=4), *Bbs4*^KO/KO^ (n=3), 2 independent experiments. Student’s t-test was used for the statistical analysis. Mid-age adult mice: *Bbs4*^+/+^ (n=7), *Bbs4*^KO/KO^ (n=5), *Bbs4*^GT/GT^ (n=3), 4 independent experiments. Kruskal-Wallis with Dunn’s Multiple Comparison Post-tests was used for the statistical analysis. Mean+SEM. **(K)** Immunoblot analysis of BBS4 expression in the brain lysates of *Bbs4*^+/+^, *Bbs4*^GT/GT^, and *Bbs4*^KO/KO^ mice. β-actin staining served as a loading control. A representative experiment out of 5 in total.

### Mouse models for studying the role of the BBSome in the immune system

Our next step was to obtain a mouse model of the BBS. We decided to use the *Bbs4*-deficient mouse for the following reasons: (I) BBS4 is an essential part of the BBSome [2], (II) *Bbs4*^KO/KO^ mouse has been shown to have a relatively severe phenotype in comparison to other BBSome-deficient mice [59, 60], (III) *Bbs4* had a relatively high expression in lymphoid tissues (Fig. 1A-B). In the following experiments, we used mice with an interrupted *Bbs4* gene with a gene-trap (GT) cassette, mice with a deletion of *Bbs4* exon 6 (KO), and mice with a *Bbs4* exon 6 flanked with LoxP sites for Cre-driven conditional deletion (Fig. 1C).

BBS4 protein was not detectable in the testes and thymi from the gene-trap *Bbs4*^GT/GT^ mice (Fig. 1D). As expected, the *Bbs4* deficiency lead to the absence of sperm flagella in testes of 30-days-old *Bbs4*^GT/GT^ males (Fig. 1E). However, we were surprised by not observing some previously reported features of BBS mouse models in the *Bbs4*^GT/GT^ mice. Whereas mating of *Bbs4*^+/GT^ heterozygotes resulted to 17% *Bbs4*^GT/GT^ pups at weaning, only 2% *Bbs4*^KO/KO^ pups were produced in mating of *Bbs4*^+/KO^ heterozygotes (Fig. 1F). This suggests that *Bbs4*^KO/KO^, but not *Bbs4*^GT/GT^, mice suffer from pre-weaning lethality with relatively strong penetrance. Moreover, *Bbs4*^KO/KO^ mice, but not *Bbs4*^GT/GT^, developed obesity (Fig. 1G-I). As expected, adult *Bbs4*^KO/KO^ mice, suffering from obesity, had elevated level of leptin in blood plasma in comparison to non-obese *Bbs4*^GT/GT^ mice, and pre-obese young *Bbs4*^KO/KO^ (Fig. 1J). Because leptin has been proposed to act as a pro-inflammatory signaling molecule, it might have an impact on the immune system of obese BBS mice.

As obesity is caused by the BBSome deficiency in the central nervous system [28], it is plausible that *Bbs4* GT allele might retain residual BBS4 expression specifically in the brain, yet we could not detect it (Fig. 1K). Another possibility is that the *Bbs4* GT allele allows for an expression of a truncated BBS4 that partially retains its function. In any case, our data suggested that the *Bbs4* GT is a hypomorphic allele, which does not lead to a complete loss of the BBS4 function.

### Alterations in the immune system of *Bbs4* deficient mice

In the next step, we analyzed the development and homeostasis of T and B cells in *Bbs4*-deficient mice. We did not observe any major alterations in the T-cell compartment in *Bbs4*^KO/KO^ and *Bbs4*^GT/GT^ mice (Fig. S1A-C). The only significant differences were decreased percentages of CD44^+^ cells among splenic CD8^+^ T cells and decreased percentage of T cells among the splenocytes of the *Bbs4*^KO/KO^ mice (Fig. S1C).

We observed an alteration of the B-cell development and/or homeostasis in *Bbs4-*deficient mice. *Bbs4*^KO/KO^, but not *Bbs4*^GT/GT^, mice showed an increased number of B220^+^ B-lineage cells in the bone marrow. Moreover, the ratio of B220^high^ and B220^low^ cells in the bone marrow was shifted towards less mature B220^low^ cells in *Bbs4*^KO/KO^ and *Bbs4*^GT/GT^ mice (Fig. 2A). Both *Bbs4*^KO/KO^ and *Bbs4*^GT/GT^ had higher percentage of IgD^−^ IgM^−^ B-cell precursors than controls, although the statistical significance was not reached (Fig. 2B). A deeper analysis of this population showed that *Bbs4*-deficiency results in a developmental block at the pre-B-cell stage, at which the pre-BCR selection occurs (Fig. 2C).

**Figure 2.**
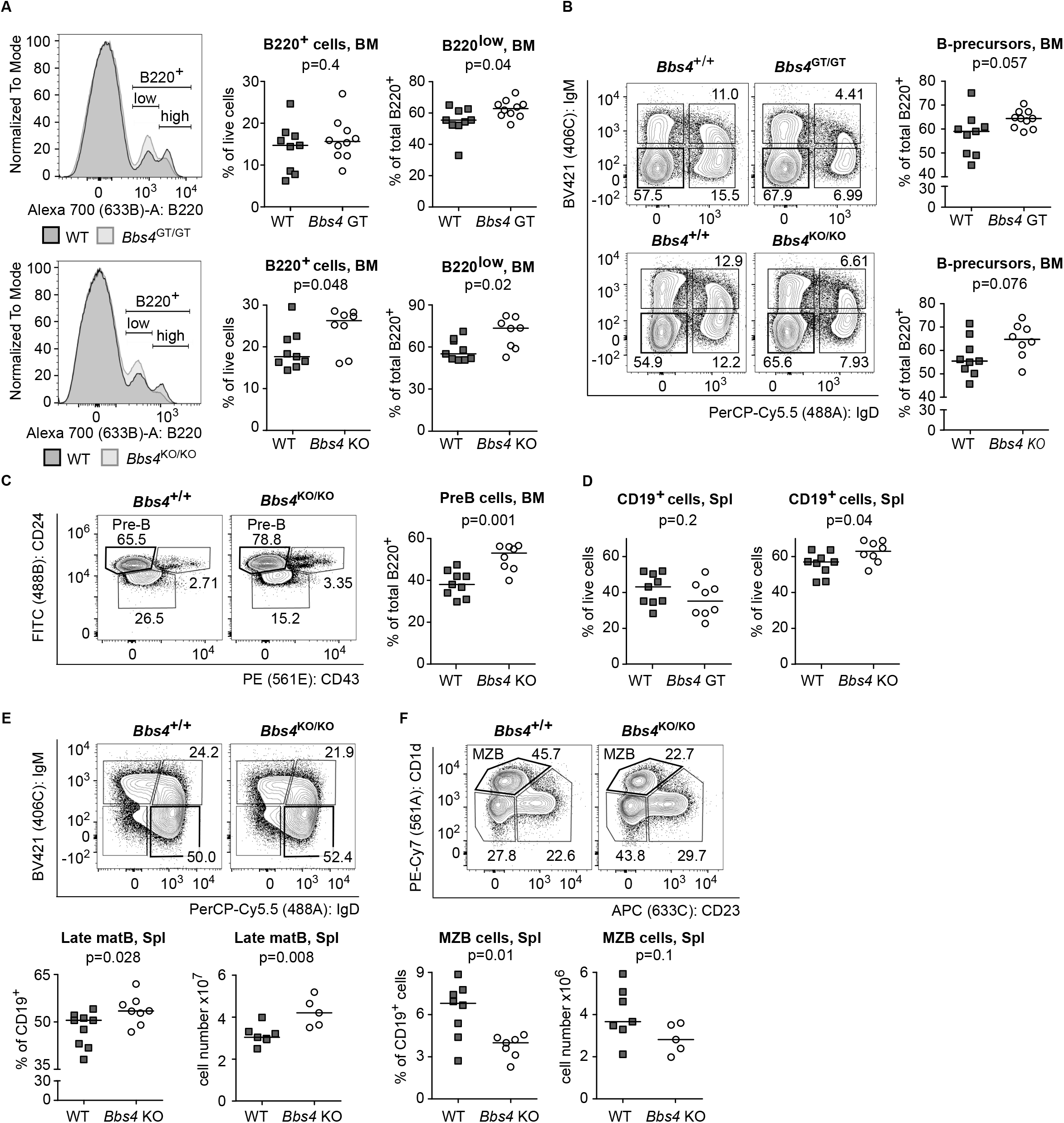
B-cell compartment is moderately affected in *Bbs4* deficient mice. **(A-F)** Cells isolated from bone marrows (A-C) and spleens (D-F) of *Bbs4*^GT/GT^, *Bbs4*^KO/KO^, and their WT littermates were analyzed by flow cytometry. **(A)** Percentages of B220^+^ and B220^low^ positive cells in bone marrow were analyzed. Representative experiments out of 6 for each strain (*Bbs4*^GT/GT^ and *Bbs4*^KO/KO^) are shown. *Bbs4*^+/+^ (n=9 per group), *Bbs4*^GT/GT^ (n=10), *Bbs4*^KO/KO^ (n=8). Unpaired t test was used for the statistical analysis (normality was checked by D’Agostino & Pearson test, p <0.05). Medians are shown. **(B)** Percentage of B-cell precursors (IgM^−^ IgD^−^) in the bone marrow. Representative experiments out of 6 for each line (*Bbs4*^GT/GT^ and *Bbs4*^KO/KO^) are shown. *Bbs4*^+/+^ (n=9 per group), *Bbs4*^GT/GT^ (n=10), *Bbs4*^KO/KO^ (n=8). Gated on viable B220^+^ cells. Unpaired t test was used for the statistical analysis (normality was checked by D’Agostino & Pearson test, p <0.05). Medians are shown. **(C)** Percentage of pre-B cells (CD43^−^ CD24^high^) in the bone marrow of *Bbs4*^KO/KO^ (n=8) and their WT littermates (n=9). A representative experiment out of 6 in total is shown. Gated on viable B220^+^ IgM^−^ IgD^−^ cells. Unpaired t test was used for the statistical analysis (normality was checked by D’Agostino & Pearson test, p <0.05). Medians are shown. **(D)** Percentage of B cells (CD19^+^) in spleen of *Bbs4*^GT/GT^ (n=8), *Bbs4*^KO/KO^ (n=8) and their WT littermates (9 mice per group) was determined. 6 independent experiments for each line were performed. Unpaired t test was used for the statistical analysis (normality was checked by D’Agostino & Pearson test, p <0.05). Medians are shown. **(E)** Percentage of late mature (IgM^−^ IgD^+^) B cells in spleen of *Bbs4*^GT/GT^, *Bbs4*^KO/KO^ (8 mice per group) and their WT littermates (9 mice per group) was determined. Representative experiments out of 6 for each strain (*Bbs4*^GT/GT^ and *Bbs4*^KO/KO^) are shown (gated on viable CD19^+^ cells). Unpaired t test was used for the statistical analysis (normality was checked by D’Agostino & Pearson test, p <0.05). Medians are shown. **(F)** Percentage of splenic MZB cells (CD23^−^ CD1d^+^) in *Bbs4*^KO/KO^ mice (n=7) and their WT littermates (n=8) was determined. A representative experiment out of 4 in total is shown (gated on viable CD19^+^, IgD^−^ IgM^+^, CD138^−^ cells). Statistical significance was calculated using two-tailed Mann-Whitney test. Medians are shown.

Interestingly, we observed increased numbers of B cells in the spleen of *Bbs4*^KO/KO^, but not *Bbs4*^GT/GT^ mice (Fig. 2D, Fig. S2A). In the spleen and lymph nodes, *Bbs4*^KO/KO^ mice have increased late mature (IgD^+^ IgM^−^) B-cell population (Fig. 2E, Fig. S2C). Furthermore, *Bbs4*^KO/KO^ showed ~2-fold decrease in the percentage of splenic marginal zone B cells (MZB) (Fig. 2F). These phenotypes were largely absent or less pronounced in the *Bbs4*^GT/GT^ animals (Fig. 2D, Fig. S2B-C), suggesting that they are caused by obesity and/or caused by the complete loss of function of BBS4 in the full KO animals.

Overall, these data show that BBSome deficiency results in a partial developmental block of B cells in the bone marrow, increase of late mature splenic B cells and relative reduction of splenic MZB cells in the periphery. The developmental block was observed in non-obese *Bbs4*^GT/GT^ mice as well, indicating that this phenotype is obesity independent.

### The role of *Bbs4* in B-cell homeostasis is not intrinsic

To investigate if the observed B-cell developmental block in *Bbs4* deficient mice is B-cell intrinsic, we generated *Bbs4*^FL/FL^ *Vav-iCre* mice with a specific deletion of *Bbs4* in the hematopoietic lineage. Surprisingly, we did not observe any signs of the altered B-cell homeostasis in *Vav-iCre* mice (Fig. 3A-B). These results suggest that the B-cell compartment in *Bbs4*^GT/GT^ and *Bbs4*^KO/KO^ mice is affected by factors extrinsic to the hematopoietic lineage, mostly likely the niche in the bone marrow and/or in the peripheral lymphoid organs.

**Figure 3.**
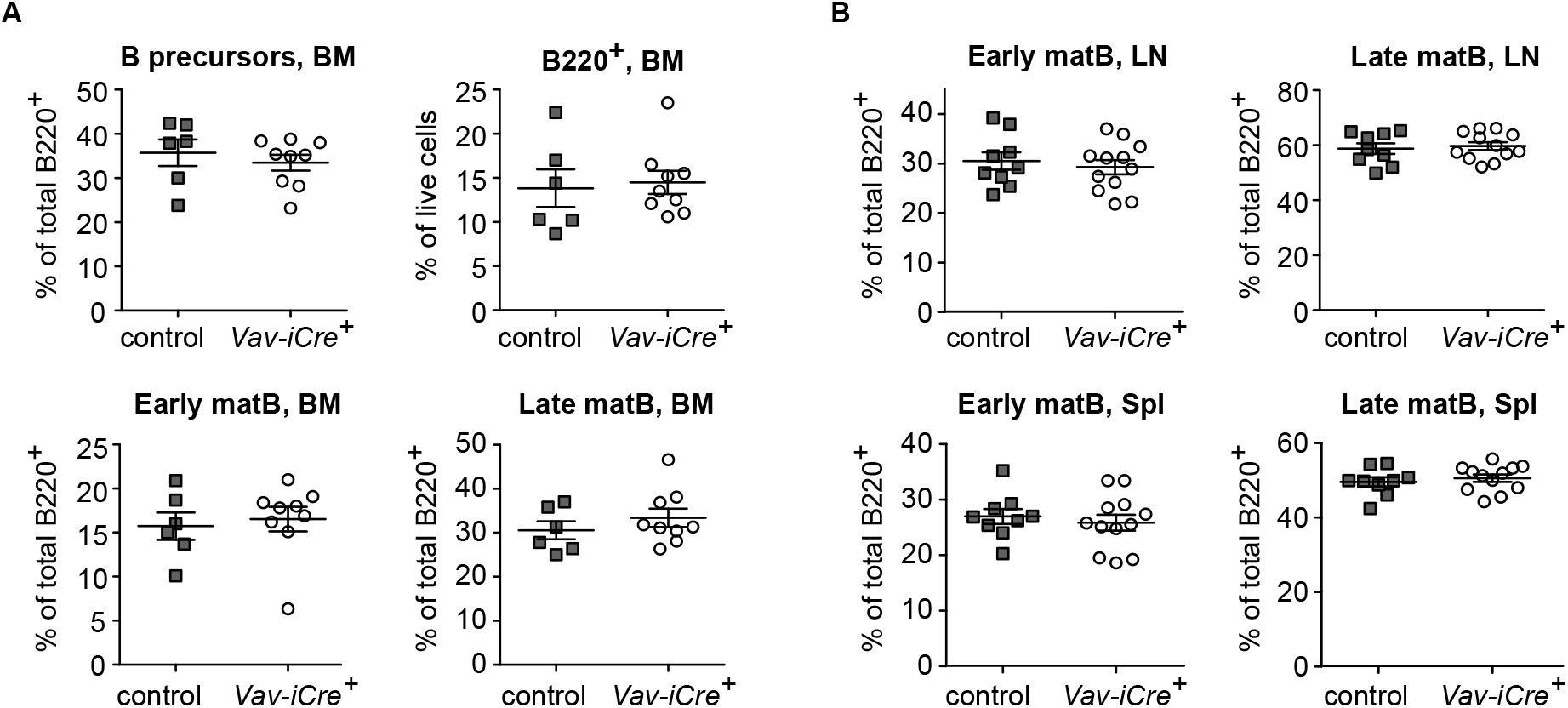
The role of *Bbs4* in B-cell development is not intrinsic. **(A-B)** Cells isolated from bone marrow (A), lymph nodes and spleen (B) of *Bbs4*^FL/FL^ *Vav-iCre*^−^ (control) and *Bbs4*^FL/FL^ *Vav-iCre*^+^ (cKO) mice were analyzed by flow cytometry. Percentages of B220-positive cells, B-cell precursors (IgM^−^ IgD^−^), early mature (IgM^+^ IgD^+^), and late mature B cells (IgM^−^, IgD^+^) were quantified. **(A)***Bbs4*^FL/FL^ *Vav-iCre*^+^ (n=8), *Bbs4*^FL/FL^ *Vav-iCre*^−^ (n=6), 4 independent experiments. **(B)***Bbs4*^FL/FL^ *Vav-iCre*^+^ (n=12), *Bbs4*^FL/FL^ *Vav-iCre*^−^ (n=9), 6 independent experiments. Mean±SEM. Statistical significance was calculated using two-tailed Mann-Whitney test, p >0.05 for all quantified parameters.

### *Bbs4* deficiency does not intrinsically influence T-cell and B-cell antigenic responses

As we observed an alteration of the B-cell homeostasis in *Bbs4*-deficient mice, we decided to investigate how it can affect the response of the adaptive immune system. First, we activated monoclonal B-cells specific to 4-hydroxy-3-nitrophenyl acetyl (NP) from B1-8 [42] and *Bbs4*^GT/GT^ B1-8 mice using NP-labeled cells and monitored the upregulation of CD69 activation marker. In this assay, we did not observe any role of *Bbs4* deficiency in the B-cell response (Fig. 4A-B). These experiments were only performed in *Bbs4*^GT/GT^ animals to exclude the possible role of obesity as an extrinsic factor.

**Figure 4.**
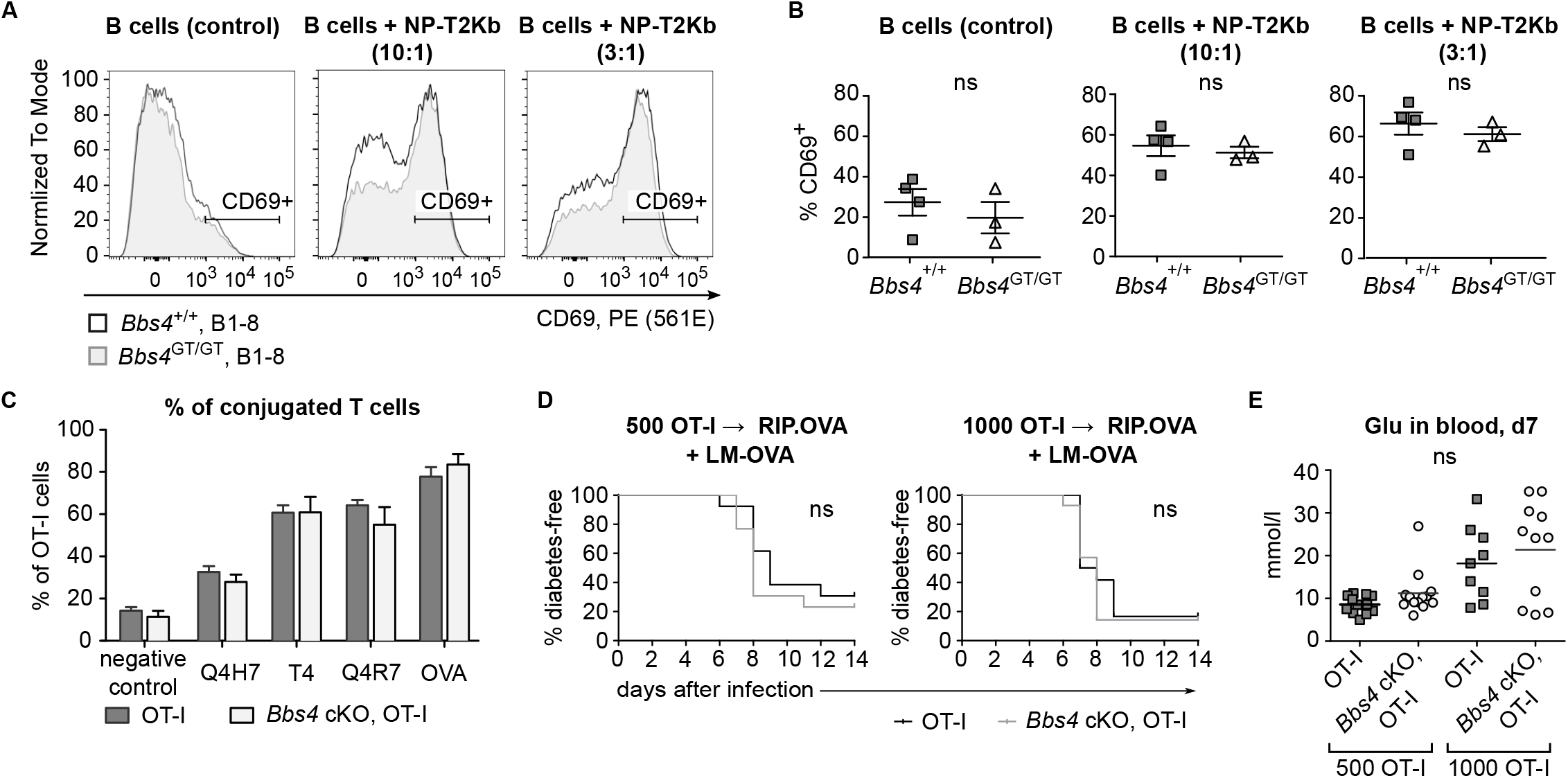
BBS4 is not required for T-cell and B-cell antigenic responses. **(A-B)** Splenocytes isolated from nitrophenyl-specific *Bbs4*^+/+^ and *Bbs4*^GT/GT^ B1-8 littermates were incubated with 4-hydroxy-3-nitrophenylacetic acid succinimide ester loaded T2-Kb cells at 37°C/CO2 incubator for 6 hours. Percentage of activated B cells (gated as CD69^+^ B220^+^ IgLλ^+^ viable cells) in the samples without T2-Kb (serve as a control), and in the samples with T2-Kb added in ratios 1:10 or 1:3 was determined by flow cytometry. (A) Representative experiment out of 3 in total. (B) 3 mice per group. Mean±SEM. Statistical significance was calculated using two-tailed Mann-Whitney test. **(C)** T cells isolated from lymph nodes of *Bbs4*^FL/FL^ and *Bbs4*^cKO^ OT-I *Rag2*^KO/KO^ littermates were loaded with CFSE and incubated with DDAO-labeled WT splenocytes loaded with OVA peptide or with the indicated altered peptide ligands at 37°C/CO_2_ incubator for 20 min. Percentage of T cells conjugated with the APCs was determined by flow cytometry. n=4 in 4 independent experiments. Mean+SEM. Statistical significance was calculated using two-tailed Mann-Whitney test, p >0.05 for all peptides. **(D-E)** 500 or 1000 T cells from *Bbs4*^FL/FL^ or *Bbs4*^cKO^ OT-I *Rag2*^KO/KO^ littermates were adoptively transferred into RIP-OVA mice followed by LM-OVA infection next day. (D) Glucose level in the urine of mice was monitored on a daily basis. The mouse was considered diabetic when it had urine glucose level ≥ 1000 mg/dL for 2 consecutive days. Statistical significance was calculated by Log-rank (Mantel-Cox) test, p >0.05 for both conditions. (E) Glucose concentration in blood on day 7 post-infection. n=12-14 animals in 4 independent experiments, mean is shown. Statistical significance was calculated using two-tailed Mann-Whitney test.

Taking into account the arising evidence that the primary cilium might share some features with the immune synapse in T cells [13, 17, 61], we expected that the T-cell functions might be compromised in the *Bbs4*-deficient mice. For this reason, we generated a *Bbs4*^FL/FL^ *Cd4-Cre* mouse line where *Bbs4* deficiency was restricted to T cells (Fig. S3A). These mice did not show any developmental alterations in the T-cell compartment (Fig. S3B-E). To study the role of BBSome in the T-cell antigenic responses, we crossed the *Bbs4*^FL/FL^ *Cd4-Cre* to TCR transgenic OT-I *Rag2*^KO/KO^ mice. This mouse generates monoclonal OT-I T cells specific for K^b^-OVA, a model antigen originating from chicken ovalbumin. We did not observe any role of BBS4 in the conjugation of OT-I T cells with antigen presenting cells loaded with OVA or with altered peptide ligands with variable affinity (Fig. 4C). Moreover, WT and *Bbs4*-deficient OT-I T cells showed the same ability to induce autoimmune diabetes upon a transfer into RIP.OVA mice expressing ovalbumin under the rat insulin promoter and subsequent priming by *Listeria monocytogenes* expressing ovalbumin [44, 52] (Fig. 4D-E). This assay examines OT-I T cells for a complex of abilities, i.e., priming by the OVA-antigen, expansion, infiltration of the pancreas, and killing the β-cells. As the onset of diabetes caused by *Bbs4*-deficient OT-I T cells was not different from the control, we concluded that BBS4 does not play an important intrinsic role in any of indicated steps of the T-cell-mediated immune response.

### BBS-induced obesity affects blood homeostasis

To investigate the possible factors predisposing BBS patients to the development of autoimmune diseases, we decided to examine the blood test results of BBS patients. Intriguingly, immunity-related parameters, such as counts of total white blood cells, neutrophils, and eosinophils, were increased in BBS patients, when compared to age and gender matched controls (BMI-random controls) (Fig. 5A). To address the role of obesity in BBS patients, we decided to use an additional set of controls with body mass indexes (BMI) matching to those of BBS patients (BMI-matched controls). We did not observe any difference when we compared the indicated leukocyte parameters between BBS patients and BMI-matched controls. In addition, we analyzed the peripheral blood of *Bbs4* deficient mice. In agreement with the analysis of the patients’ blood tests, we did not observe major differences between WT and non-obese *Bbs4*^GT/GT^ mice. However, obese *Bbs4*^KO/KO^ mice showed higher total white blood cell count than WT controls (Fig. 5B) in line with the data from patients (Fig. 5A). These results indicate that obesity in BBS patients and in *Bbs4*-deficient mice has an impact on the leukocyte homeostasis. Moreover, BBS patients showed significantly higher C-reactive protein (CRP) levels than BMI-matched and BMI-random controls (Fig. 5C), which indicates that obesity is not the only factor influencing the immune system of BBS patients.

**Figure 5.**
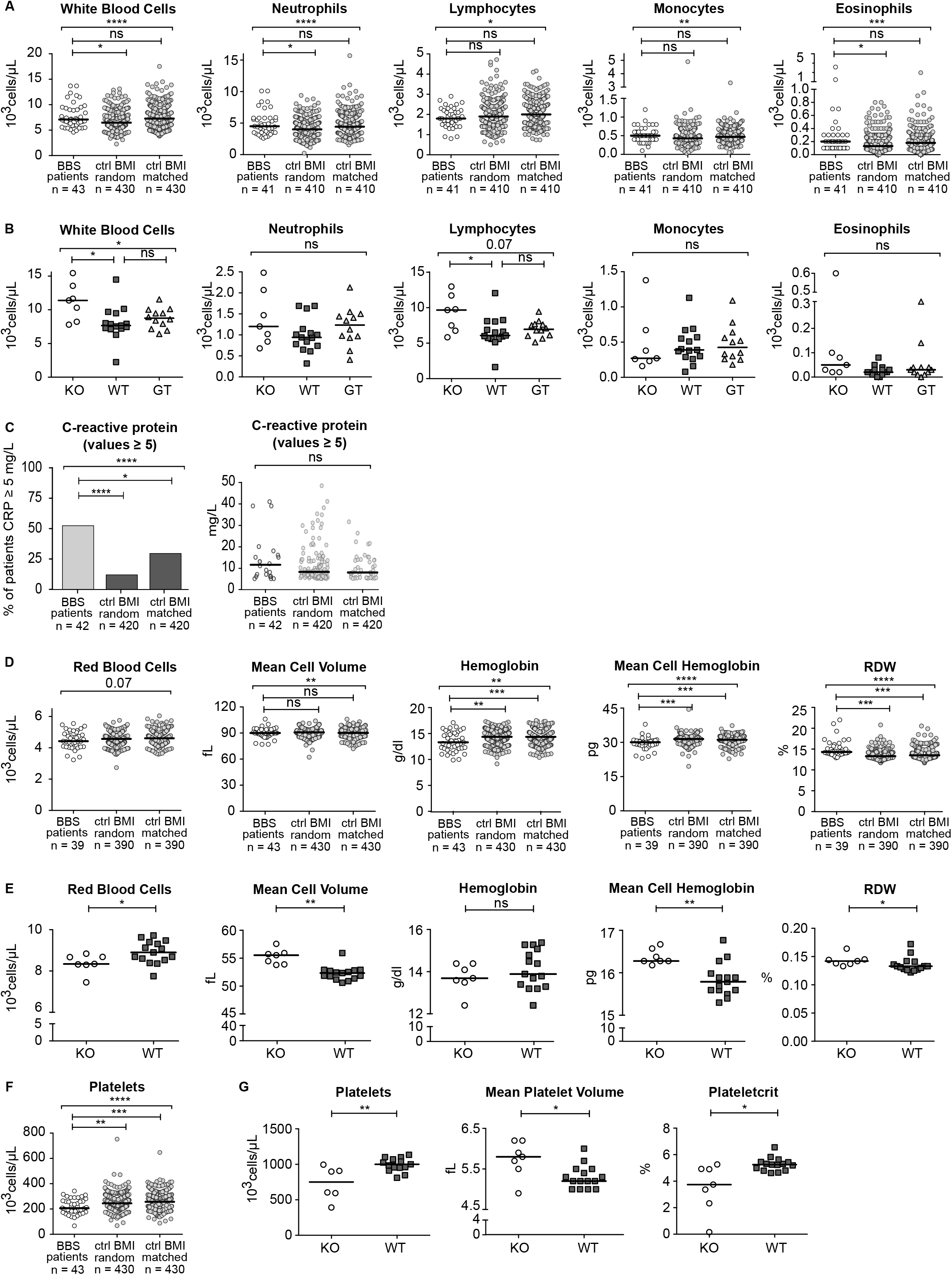
Blood homeostasis is altered in BBS patients and *Bbs4* deficient mice. **(A, C, D, F)** Results of the blood tests of BBS patients from the Guy’s Hospital and Great Ormond Street Hospital were extracted from the medical records and compared to two sets of healthy controls obtained from the UK Biobank. The BMI-random controls were age- and gender-matched to the set of BBS patients, with random BMIs. The BMI-matched controls were matched for age, gender, and BMI. Both data sets of healthy controls were selected as 10-fold larger than the set of BBS patients to obtain higher statistical power. Median is shown. Kruskal-Wallis test was used for the statistical analysis in A, D and F. In C, the percentages of patients or controls having CRP≥5 were compared using Fisher’s exact test with post-hoc Sidak correction for multiple comparisons. **(B, E, G)** Indicated parameters were measured using the blood from 20-21 weeks old *Bbs4*^KO/KO^ (n=7), *Bbs4*^GT/GT^ (n=12), and *Bbs4*^+/+^ (n=15) mice. Kruskal-Wallis test with Dunn’s Multiple Comparison Post-tests was used for the statistical analysis. Median is shown.

Notably, the homeostasis of red blood cells was altered in BBS patients as well as in the *Bbs4*^KO/KO^ mice, although at different levels (Fig. 5D-E). BBS patients showed low overall hemoglobin levels caused by a mild decrease in the red blood cell count and low red blood cell hemoglobin (Fig. 5D), indicating a possibility of a reduced oxygen transport capacity. Interestingly, the comparison of BBS patients with BMI-matched controls showed that the alteration of the erythroid compartment was not caused by obesity. The mouse model of BBS showed a decreased number of red blood cells, which was compensated by enlarged red blood cell volume (Fig. 5E).

In addition, we observed decreased platelet counts both in BBS patients and *Bbs4*^KO/KO^ mice (Fig. 5F-G). The reduction of platelets in BBS patients was not obesity-dependent. Furthermore, *Bbs4*^KO/KO^ mice showed higher mean platelet volume and lower platelecrit than WT mice (Fig. 5F-G), indicating enhanced removal of platelets in the periphery.

Altogether, our results suggest the role of the BBSome in the immune tolerance, hematopoiesis and/or blood homeostasis. Most of the effects seem to be extrinsic to the hematopoietic compartment as revealed by using tissue-specific knock-out mouse model and comparison of patients to BMI-matched controls. However, altered CRP levels, red blood cell, and platelet homeostasis in BBS patients are obesity-independent.

## Discussion

In this work, we have shown that the BBS is connected with changes in the immune system and blood homeostasis and with higher incidence of autoimmune diseases. First, *Bbs4* depletion affects B-cell development and red blood cell and platelet homeostasis, and second, it leads to obesity, which induces changes in the immune populations.

Lymphocytes are usually considered as a classical example of non-ciliated cells [10, 11], although some contradicting studies were published [62, 63]. Importantly, T cells repurpose a number of proteins associated with the ciliary transport for the formation of the immunological synapse. Three prominent signaling molecules in T cells (TCR, LAT, LCK) are transported to the immunological synapse via the intraflagellar transport machinery (IFT) [15, 17, 61]. As the BBSome directly interacts with the IFT [64–66], we hypothesized that the BBSome might have a cilia-independent role in the T-cell activation. However, our data do not support this hypothesis, because the immune response of *Bbs4*-deficient T cells was not affected.

Some ciliary proteins, including BBSome-interacting partner IFT20, and SHH signaling components, play a role in the early T-cell development. It has been shown that the *Lck*-*Cre* driven ablation of *Ift20* in early stages of T-cell development impairs the maturation of thymocytes, leading to the reduction of peripheral T cells [67]. Interestingly, *Ift20* ablation at later stages of T-cell development using *Cd4*-*Cre* had only mild effect on the T-cell maturation [15, 67]. Similar findings were shown using *Lck-Cre* and *Cd4*-*Cre* driven ablation of SMO, a receptor involved in SHH signaling [24, 68]. In our study, the T-cell lineage was largely unaffected in the *Bbs4*^KO/KO^ mouse and in *Cd4*-*Cre* driven *Bbs4*^cKO/cKO^ mice, which suggests that the BBSome is dispensable for SHH signaling in thymocytes and for T-cell development in general.

In contrast to the T-cell lineage, we did observe alterations in the B-cell compartment of *Bbs4*-deficient mice. *Bbs4*^KO/KO^ and hypomorphic *Bbs4*^GT/GT^ mice showed a partial developmental arrest at the pre-B cell stage in the bone marrow, which was not caused by obesity. Specific ablation of *Bbs4* in hematopoietic cells did not impair the development of B cells indicating that other extrinsic factors control B-cell development in the bone marrow. Bone marrow niches include different cells of non-hematopoietic origin, which have been shown to regulate the hematopoietic cell development and recirculation, including reticular stromal cells, perivascular cells, osteolineage cells, and non-myelinating Schwann cells [69, 70]. Therefore, it is possible that the impeded B-cell development in the BBS mouse model is a result of the altered microenvironment in the bone marrow.

Moreover, *Bbs4*^KO/KO^ mice show high frequency and absolute numbers of IgM^+^ IgD^+^ late mature B cells, whereas the frequency of MZB cells was decreased in the spleen. MZB cells are known as the main producers of IgM antibodies [71] which play a role in the early line of the immune defense. The changes in the peripheral B-cell compartment were statistically significant only in *Bbs4*^KO/KO^, but not in *Bbs4*^GT/GT^, pointing to the hypomorphic nature of the *Bbs4*^GT^ allele and/or to the role of obesity in the B-cell homeostasis.

*Bbs4*^KO/KO^ mice, similarly to the majority of BBS patients, develop obesity which influences the immune system. Obesity can induce the state of low-grade metabolic inflammation [72], characterized by elevated TNFα, IL-6, and CRP in blood [73–76] and adipose tissue [77]. Obesity is a risk factor for some autoimmune disorders, including multiple sclerosis [78–80], systemic lupus erythematosus [81], rheumatoid arthritis [82–85], and autoimmune diabetes mellitus [86, 87]. The risk of Raynaud’s and celiac diseases is decreased in obesity [86], and the risk of inflammatory bowel disease [88–90] and hypothyroidism [91–93] remains controversial. In this work, we have shown elevated incidence of certain autoimmune disorders in BBS cohorts, as well as altered composition of blood cells in BBS patients and BBS mouse models. The comparison of BBS patients and non-BBS obese controls showed that the alteration of the white blood cell count in BBS patients is caused by obesity, which was supported by the murine data. Altogether, our results suggest that obesity might substantially contribute to the high incidence of certain autoimmune diseases in BBS patients. However, the data also indicate that obesity is not the only factor altering the immune status of BBS patients. This is showed for example by the increased CRP level in BBS patients, which can not be fully explained by the high BMI. It should be noted that BBS patients are under various medications, some of which can potentially alter the blood homeostasis.

One of the major players in obesity-associated inflammation is leptin, an adipocyte-derived hormone which acts as a pro-inflammatory cytokine [94, 95]. Leptin receptor is expressed by different cell types including T cells, in which it promotes T-cell activation and proliferation. [32, 96]. It has been shown that the BBSome is required for the cell surface delivery of the leptin receptor in human kidney epithelial cell line HEK293 [28]. However, we did not observe immune response defects in murine T cells deficient in *Bbs4*, implying that the BBSome is not required for the proper function of the leptin receptor in T cells.

In addition, we observed thrombocytopenia both in BBS patients and in *Bbs4*-null mice; and furthermore, platelets from these mice had larger volume. There are two major causes of thrombocytopenia, the first one is a decreased production of platelets in the bone marrow, and the second is an over-destruction of platelets on the periphery [97]. Since the size of the immature platelets is larger than that of the mature ones, large platelet volume combined with thrombocytopenia indicates active platelet production in the bone marrow compensating for their loss in the periphery [98]. This state can be a result of immune-mediated processes such as immune thrombocytopenia; it is also seen in disseminated intravascular coagulopathy, thrombotic thrombocytopenic purpura, and sepsis [99, 100]. Unfortunately, we do not have results of mean platelet volume from BBS patients, but we can presume that the reasons of thrombocytopenia in BBS patients and mice are similar, suggesting a role of the peripheral destruction of platelets.

To our knowledge, the immune system in ciliopathies has not been investigated yet, although it can be affected in multiple ways. First, obesity is a common feature of ciliopathies such as BBS and Alström syndrome [101]. Second, the deficiency of ciliary proteins might have different effects on immunity, including immunological synapse defects, which needs to be further elucidated. Third, non-hematopoietic ciliated cells interact with leukocytes and play a role in the immune defense. Here, we show that the absence of a ciliary protein BBS4 affects the immune system of humans and mice in two ways, i.e., obesity-dependent (changes in leukocyte homeostasis), and obesity-independent (high CRP levels in BBS patients, partial developmental block in the B-cell lineage).

## Acknowledgement

We thank Ladislav Cupak for technical assistance and genotyping of mice and Dr. Alena Moudra for the assistance with *Listeria* infection. This study was supported by the Czech Science Foundation (17-20613Y to MH), and the Charles University Grant Agency (1706119 to OT), and the National Institute for Health Research Biomedical Research Centre at Great Ormond Street Hospital for Children NHS Foundation Trust and University College London. The Group of Adaptive Immunity is supported by an EMBO Installation Grant (3259 to OS) and the Institute of Molecular Genetics of the Czech Academy of Sciences core funding (RVO 68378050). This research has been conducted using the UK Biobank Resource under application number #40103. OS is supported by the Purkyne Fellowship provided by the Czech Academy of Sciences. OT, VN and AP are students partially supported by the Faculty of Science, Charles University, Prague.

The animal facility of the IMG is a part of the Czech Centre for Phenogenomics and the work there was supported in part by following grants: LM2015040, OP RDI CZ.1.05/2.1.00/19.0395, OP RDI BIOCEV CZ.1.05/1.1.00/02.0109 provided by the Czech Ministry of Education, Youth and Sports and the European Regional Development Fund.

## Author contribution

MH and OS conceived the study and were in charge of the overall direction and planning. OT, VN, AD, AP, and OS performed the animal experiments. EF, KS, and PB collected the data of BBS patients from the Great Ormond Street Hospital and Guy’s Hospital in London. VN filtered and analysed the patients’ data. ZT contributed to the histological analysis of murine testes. OT, VN, and OS wrote the manuscript with the contribution of all the other authors.

